# Mechanisms of viral budding through cellular membranes

**DOI:** 10.64898/2026.06.10.731270

**Authors:** Shuyang Zhang, Siyu Li, Miguel Angel Coronado-Ipiña, Mauricio Comas-Garcia, Ajay Gopinathan, Paul van der Schoot, Roya Zandi

**Affiliations:** Department of Physics and Astronomy, University of California Riverside, Riverside, CA 92521, United States; Department of Physics and Astronomy, California State Polytechnic University, Pomona, CA 91768, United States; Department of Sciences, Autonomous University of San Luis Potosi, Mexico; Research Center for Health Sciences and Biomedicine, Autonomous University of San Luis Potosi; Department of Physics, University of California, Merced, California 95340, United States; Department of Applied Physics and Science Education, Eindhoven University of Technology, P.O. Box 513, 5600 MB Eindhoven, The Netherlands

## Abstract

Budding is a fundamental membrane-remodeling process central to many cellular functions and is exploited by numerous enveloped viruses to acquire their lipid envelopes. Despite extensive molecular characterization, the physical mechanisms that determine whether budding proceeds to completion or becomes stalled remain unclear. Here, we develop a theoretical model based on the Helfrich elastic formalism to investigate how membrane geometry and boundary conditions regulate the elastic energy of viral budding. We analyze two representative cases: budding from a flat membrane, characteristic of HIV-1 and alphaviruses, and budding from a vesicle, as observed for SARS-CoV-2 in the ER–Golgi intermediate compartment (ERGIC). Our results reveal distinct energetic pathways: vesicle-like geometries exhibit a stronger energetic bias toward closure, whereas flat membranes develop extended low-slope regions in the energy landscape that can hinder completion. Relaxing far-field boundary constraints reduces the energetic cost associated with membrane area conservation and renders the flat-membrane case energetically comparable to the vesicle case, providing a physical explanation for why viruses frequently bud adjacent to one another or within pre-curved membrane regions. Comparison with thin-section TEM images of alphavirus budding shows results consistent with the theoretical membrane profiles. Together, these findings establish how curvature coupling, boundary flexibility, and local membrane geometry cooperate to control the efficiency and completion of membrane budding.

## I. INTRODUCTION

Budding is a critical step in numerous cellular processes, including endocytosis, exocytosis, the formation of multivesicular bodies, retrograde trafficking, exosome release, and material exchange between cells. Many enveloped viruses also exploit budding to acquire their membranes. In these cases, interactions between the preassembled nucleocapsid core and the host plasma membrane lead to the encapsulation of the nucleocapsid by the lipid bilayer, as seen in alphaviruses^1,2^. In contrast, in the human immunodeficiency virus (HIV-1), assembly and budding are tightly coupled: the Gag proteins that form the immature virion assemble and bud simultaneously through the plasma membrane^3–5^.

Budding can occur in various cellular compartments, including the endoplasmic reticulum (ER), the Golgi apparatus, and the ER–Golgi intermediate compartment (ERGIC). For instance, clathrin and caveolae assemble at the plasma membrane to produce endocytic vesicles^6–9^, whereas COPI and COPII coat proteins bud from Golgi and ER membranes^10,11^. Coronaviruses such as Severe Acute Respiratory Syndrome Coronavirus 2 (SARS-CoV-2) employ an intracellular budding pathway^12–14^. SARS-CoV-2 forms infectious virions through the coordinated assembly of its structural proteins and genomic RNA, which bud from the lipid membranes of the ERGIC. The virus encodes four structural proteins—nucleocapsid (N), membrane (M), envelope (E), and spike (S)—whose interactions drive budding and virion morphogenesis. Studies by Newman *et al*.^15^ suggest that the M proteins form a lattice-like scaffold that induces membrane curvature and drives budding, while the N and E proteins modulate this process, although their specific roles remain incompletely understood.

Recent coarse-grained molecular dynamics simulations have provided deeper insight into the molecular mechanisms governing coronavirus budding. These simulations demonstrate that M proteins aggregate and induce curvature, enabling budding through a spherical vesicle that mimics the ERGIC^16^. In contrast, simulations performed on flat membranes under otherwise identical conditions show that budding fails to occur unless the protein–protein interactions are strong enough to overcome the absence of pre-existing curvature^17–20^. This observation raises a fundamental question: why is budding more efficient through vesicle-like structures such as the ERGIC than through flat membranes?

Fig. 1 illustrates these two distinct budding pathways. The Fig. 1(a) shows immature HIV-1 particles budding outward through the plasma membrane^3^, while the Fig. 1(b)-(d) depicts distinct aspects of coronavirus budding through the curved membranes of the ERGIC^13^. These images highlight the contrasting geometries that underlie the two mechanisms and motivate our theoretical analysis of how membrane curvature and boundary conditions influence the energetics of budding.

**FIG. 1.**
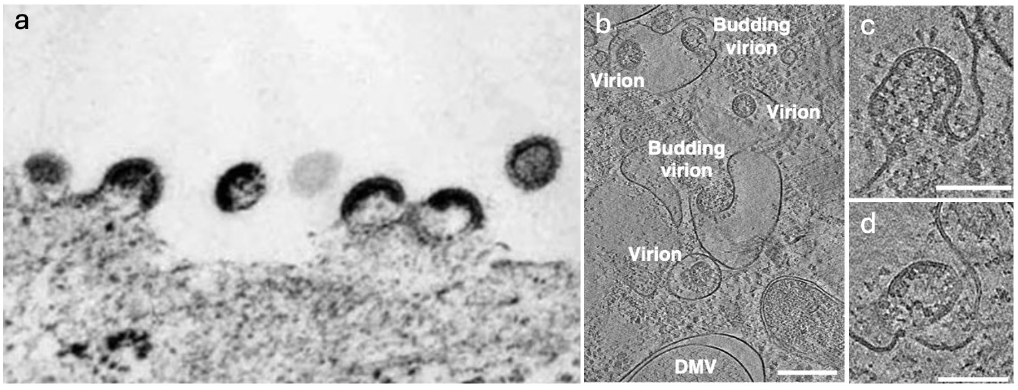
Images of HIV and coronavirus budding. (a) Immature HIV particles budding through the plasma membrane (adapted from Ref.^3^). (b–d) Coronavirus budding through the ERGIC (adapted from Ref.^13^, licensed under CC BY 4.0). (b) Overview of budding events at the ERGIC membrane and intracellular virions released into the ERGIC lumen. (c,d) Early virion budding stages, showing S proteins and vRNPs accumulated at the luminal and cytosolic faces of the ERGIC membrane, respectively. Scale bars: (b) 200 nm; (c–d) 100 nm.

Despite its biological importance, the physical principles that govern the initiation, progression, and completion or stalling of budding remain poorly understood. To address this question, we developed a theoretical model to investigate how membrane geometry and boundary conditions influence the energetics and mechanics of budding. Specifically, we employed the Helfrich formalism to describe membrane elasticity and solved the corresponding shape equations to determine the membrane configurations and energies along the budding pathway^21–25^. For simplicity, we consider a regime in which a protein-enriched cap is already present and fix the cap area, allowing us to decouple equilibrium membrane geometry from protein diffusion and assembly kinetics, which lie beyond the scope of this work.

Our analysis focuses on two representative geometries: budding from a flat membrane and budding from a vesicle^26,27^. These two cases mimic, respectively, the plasma-membrane budding of viruses such as HIV-1 and alphaviruses^28^, and the vesicular budding pathway of SARS-CoV-2 through the ERGIC. By systematically comparing these configurations, we identify how bending rigidity, the membrane area constraint, and boundary conditions control the shape evolution and the underlying energy landscape of the budding process. This comparison reveals distinct energetic pathways: budding from vesicles exhibits a steeper energy slope toward bud closure, whereas budding from flat membranes can develop extended regions with a small energy gradient, which hinder completion.

Cryo-EM images of HIV-1 and thin-section TEM images of alphaviruses show that viral budding rarely occurs on perfectly flat membranes. Instead, it often initiates in regions already deformed by neighboring buds or underlying curvature. These experimental observations suggest that local membrane deformation reduces the energetic barrier for bud completion and promotes cooperative budding. To test this prediction quantitatively, we compared the theoretical membrane shapes with thin-section TEM images of alphavirus budding. The predicted contours closely matched the experimental membrane geometry, accurately capturing both the cap curvature and neck morphology. Minor deviations on one side of the bud reflected intrinsic membrane asymmetry, which we reproduced by introducing a slightly asymmetric boundary condition. This agreement validates the predictive power of the model and highlights the importance of local curvature and boundary flexibility in shaping the membrane during viral assembly.

Together, these findings provide a unified physical picture of viral budding. Local curvature and boundary constraints determine whether budding proceeds to completion or becomes stalled. Vesicle-like geometries, such as those of the ERGIC, naturally promote efficient closure due to their inherent curvature, whereas flat membranes require boundary relaxation or assistance from neighboring buds to complete budding. This cooperative, curvature-assisted mechanism offers a physical explanation for the spatial clustering of budding sites observed in many enveloped viruses.

## II. METHODS

### A. Theory

To investigate the energetics of the virus–membrane complex during budding, we consider two representative geometries: (i) a flat membrane and (ii) a spherical vesicle-shaped membrane. In both cases, membrane-associated proteins assemble on the surface to form a two-dimensional patch, referred to as the cap region. Even though the mature HIV capsid adopts a conical shape^29^, the immature HIV particle is spherical and consists of Gag proteins assembling on the plasma membrane^30^. In contrast, for coronaviruses, the cap corresponds to M proteins assembling on a vesicle-shaped surface that mimics the ERGIC^31^. Because the size of a typical virus is much smaller than the radius of the host cell, the plasma membrane can be locally approximated as flat in the HIV case. Electron microscopy images of budding regions at various growth stages^32^ reveal that the cap regions adopt a nearly spherical shape characterized by an approximately constant curvature.

Guided by these observations, we follow the approach of Forêt^33^, assuming that the cap region maintains a spherical-cap geometry with uniform spontaneous curvature. The cap therefore imposes its preferred curvature on the portion of the membrane to which it adheres, locally deforming the underlying lipid bilayer. This curvature defines the geometry of the budding region and serves as a key parameter in the energetic analysis of the membrane shape. The extent of this deformation is quantified by the angle *α*, defined as the angle between the tangent to the cap boundary and the *r*-axis, as illustrated in Fig. 2.

**FIG. 2.**
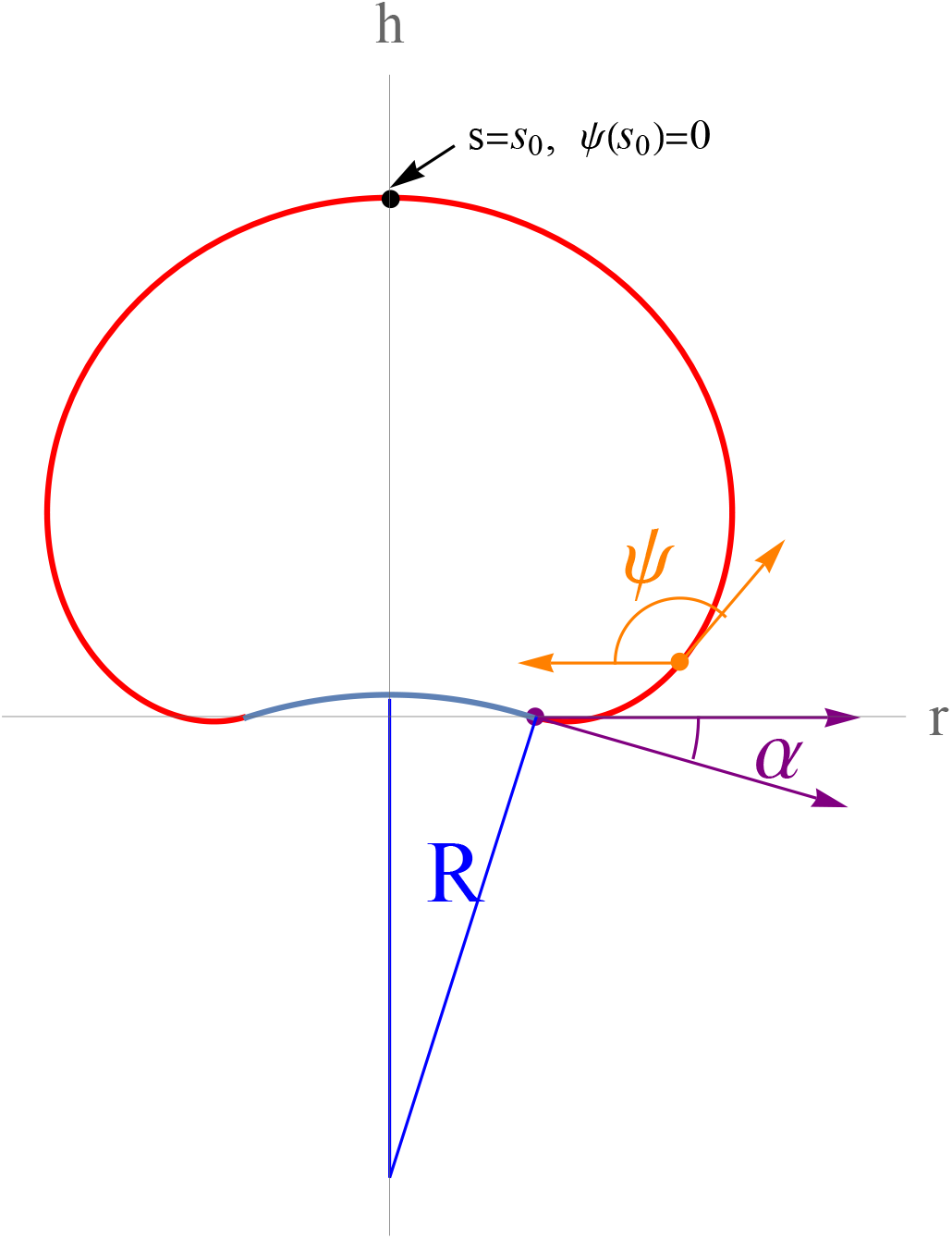
Schematic illustration of the vesicle budding geometry. The membrane surface is modeled as an axisymmetric contour described by the arc length *s* (red line), measured from the boundary of the protein-covered cap region (blue). The cap is modeled as a spherical segment of radius *R* extending up to an opening angle *α*, defined as the angle between the tangent to the membrane at the cap boundary and the radial axis. The local membrane configuration is parameterized by *r*(*s*), the radial distance from the symmetry axis (*h*-axis), *h*(*s*), the height along this axis, and *ψ*(*s*), the angle between the tangent to the membrane contour and the negative radial direction. As defined in the main text, the membrane shape satisfies 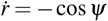 and 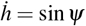. The total contour length is denoted by *s*_0_.

The initial flat configuration of the cap region corresponds to *α* = 0, whereas *α* = *π* represents a fully closed vesicle. Assuming that budding begins once a sufficient number of M proteins have aggregated, and that their number remains constant throughout SARS-CoV-2 budding process, the cap area is taken to be fixed, while its radius *R* gradually decreases as budding proceeds, approaching the spontaneous radius of curvature *R*_0_ preferred by the embedded proteins.

This geometric relationship captures the morphological evolution of the membrane as it deforms under the curvature imposed by the embedded patch of M proteins, providing the basis for calculating the bending energy throughout the budding process.

Following the work of Forêt and Deserno^33,34^, we calculate the membrane shape and elastic energy using the standard formalism of fluid membrane elasticity under axisymmetric geometry. Fig. 2 shows a schematic of the vesicle budding geometry. The *h*-axis defines the symmetry axis of the system, and the contour of the membrane is parameterized by the arc length *s*, measured from the boundary of the cap region. At each point along the contour, the membrane geometry is described by three variables: (i) *r*(*s*), the radial distance from the symmetry axis; (ii) *h*(*s*), the coordinate along the *h*-axis; and (iii) *ψ*(*s*), the angle between the tangent to the contour and the negative radial direction. Together, these variables define the local membrane shape and provide the basis for evaluating the curvature, surface area, and elastic energy in the subsequent analysis. We note that

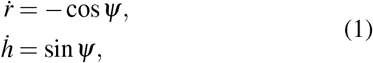

where the dot denotes differentiation with respect to the arc length *s*.

In the standard formalism of fluid membrane elasticity, the total energy contains two principal contributions: the bending energy and a surface “tension” term that enforces an area constraint. The corresponding expressions are

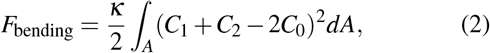

and

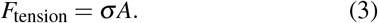

Here, *F*_bending_ follows the standard Helfrich formulation^21^, where *κ* denotes the bending rigidity of the membrane, *C*_1_ and *C*_2_ are the principal curvatures, and *C*_0_ is the spontaneous curvature. In Eq. 3, *σ* acts as a Lagrange multiplier enforcing the constraint of constant total membrane area *A*^35,36^. Since the cap shape is fixed for each value of *α*, line tension does not enter into the minimization that determines the membrane contour. Its energetic contribution is therefore not included here and will be discussed in the Results section.

Each energy term is separated into contributions from the protein-covered region and the surrounding free membrane. In the cap region, where the membrane adopts a spherical-cap geometry with constant curvature, the bending energy, *F*_cbend_ can be evaluated analytically. Because the cap region represents a densely packed protein shell, it can be treated as rigid. Consequently, the contribution from Eq. 3 to the total energy within the cap region is neglected. The remaining portion of the membrane, which is free of proteins, contributes additional elastic energy that is obtained from the numerical solution of the shape equations.

The bending energy of the cap can be written as

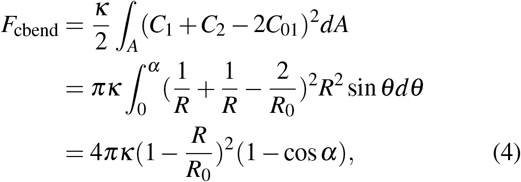

where *R* is the radius of curvature of the cap, given by 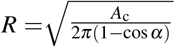 and *A*_c_ = *f A* denotes the area of the cap, where represents the fraction of the total area *A*. The principal curvatures in the spherical-cap geometry are *C*_1_ = *C*_2_ = 1*/R*. The quantity 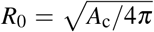 represents the radius of curvature when the budding process is complete, corresponding to a fully closed spherical vesicle. The spontaneous curvature of the cap is thus given by *C*_01_ = 1*/R*_0_. Here *θ* denotes the polar angle in spherical coordinates.

In the region of the membrane not covered by proteins, the local elastic energy is determined by the functions *r*(*s*) and *ψ*(*s*). The bending energy of the protein-free membrane region, *F*_fbend_, can be written as

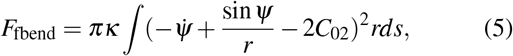

which represents an integral over local bending contributions and thus constitutes a functional of the membrane shape. The parameter *C*_02_ = 0 reflects the tendency of the protein-free membrane region to remain flat. Using the rotational symmetric arc-length parametrization shown in Fig. 2, the two principal curvatures are given by (sin *ψ*)*/r* and 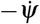, where the dot denotes differentiation with respect to the arc length *s*.

The energy functional can hence also be expressed as *É* = ∫ *Lds*, in terms of the Lagrangian *L* given by

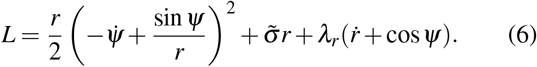

The first term in the Lagrangian represents the bending contribution, the second enforces an area constraint through the Lagrange multiplier, and the last term imposes the non-holonomic geometric constraint 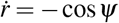 via the Lagrange multiplier *λ*_*r*_(*s*). Note that a non-holonomic constraint involves derivatives of the coordinates rather than the coordinates themselves; here, it links *r*(*s*) and its derivative through the local tilt angle *ψ*(*s*), ensuring a continuous and properly parameterized membrane contour along its arc length. The numerical factors of 2*π* (arising from the azimuthal integration in the axisymmetric geometry) have been absorbed into the definition of the Lagrangian for later convenience. All lengths are expressed in units of a characteristic scaling length *a* = 1 *nm*. Energies are measured in units of the bending rigidity *κ*, leading to the definition 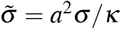. The factor *a*^2^ is introduced to render 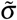 dimensionless, since *σ* has units of energy per unit area. We emphasize that the associated tension-like term 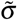 appears as a Lagrange multiplier in the variational formulation and explicitly enforces the constraint of constant total membrane area in the governing equations.

Taking the SARS-CoV-2 virus as an example, we set *R*_0_*/a* = 40. This choice allows the model to be easily rescaled for viruses of different size. No volume constraint is applied for either vesicle or planar geometries, since the high permeability of the membrane allows rapid water exchange on the characteristic timescale for budding, ensuring osmotic equilibration across the bilayer. Stretching contributions are neglected in our description, as bending is energetically far more favorable than stretching during membrane deformation^37^.

Since the Lagrangian *L* does not explicitly depend on the arc length *s*, the corresponding Hamiltonian is conserved. This follows directly from the translational invariance of the system along *s*: if the independent variable does not appear explicitly in the Lagrangian, Noether’s theorem guarantees the existence of a conserved quantity^34^, which in this case is the Hamiltonian *H*. Moreover, numerical integration is most conveniently performed for systems expressed as first-order differential equations. These considerations motivate the adoption of a Hamiltonian formulation. Defining the conjugate momenta as

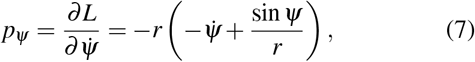

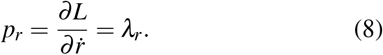

We subsequently perform a Legendre transform to obtain the Hamiltonian, from which the shape equations follow:

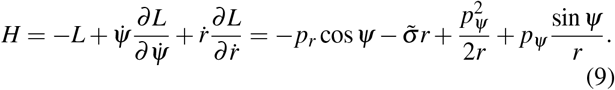

The resulting first-order differential equations that describe the equilibrium shape of the membrane are:

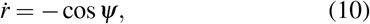

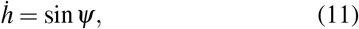

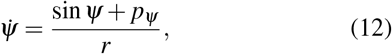

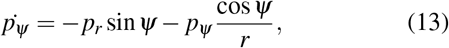

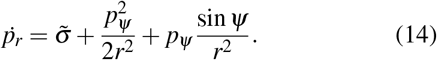

Eqs. (10) and (11) establish the geometric relations among r, h and *ψ*, follow directly from Eq. (1). Eq. (11) is a geometric relation that follows from the arc-length parameterization of the membrane contour and is used to reconstruct the membrane height profile. Eq. (12) is obtained by rearranging Eq. (7). Eqs. (13)–(14) are derived from the minimization of the energy functional 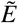. Together, Eqs. (10)–(14) form a complete set that determines the membrane shape at mechanical equilibrium.

The equations need to be supplemented by boundary conditions. The first three ensure continuity at the interface between the free and cap regions at *s* = 0:

- *r*(0) = *R* sin *α* enforces geometric continuity of the radial coordinate at the junction between the cap and free membrane;
- *h*(0) = 0 defines the vertical origin of the coordinate system at this interface;
- *ψ*(0) = *π* + *α* ensures continuity of the membrane, corresponding to the contact angle between the cap and the adjoining membrane.

We impose the boundary condition 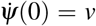. The boundary curvature 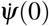 is not specified a priori but is determined numerically using a shooting method that enforces the farfield boundary conditions, at *s* = *s*_0_. Because the contact angle is prescribed, the usual torque-balance condition cannot be imposed at the boundary. The correct curvature therefore emerges from the numerical solution that satisfies the far-field constraints. From Eq. (7), together with the expressions for *r*(0) and *ψ*(0), it follows that

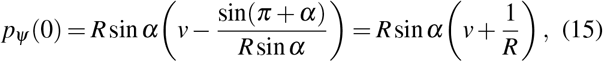

where the identity sin(*π* + *α*) = − sin *α* has been applied. Combining the Hamiltonian expression in Eq. (9) with the result for *p*_*ψ*_ (0) in Eq. (15), the initial value of *p*_*r*_(0) can be written explicitly as

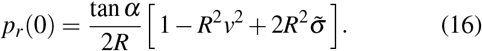

To ensure proper vesicle closure at the total membrane arc length *s* = *s*_0_, the following boundary conditions are imposed:

- *r*(*s*_0_) = 0, enforcing contour closure along the symmetry axis;
- *ψ*(*s*_0_) = 0, ensuring the membrane is tangent to the axis at the closure point and eliminating curvature discontinuities.

The set of coupled first-order differential equations is solved numerically using a fourth-order Runge–Kutta integration scheme. A shooting method is employed to determine the appropriate values of the initial condition 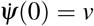 such that all boundary conditions are simultaneously satisfied. The value of *v* must be determined with high precision to satisfy the boundary conditions. Integration proceeds from the cap–membrane interface (*s* = 0) toward the symmetry axis (*s* = *s*_0_). The algorithm iteratively adjusts the initial conditions until *r*(*s*_0_) = 0 and *ψ*(*s*_0_) = 0 are met within a prescribed tolerance (typically 0.005).

To ensure numerical stability and generality, all quantities are expressed using the characteristic scaling length *a* and bending rigidity *κ*. Step sizes Δ*s* are chosen adaptively to control local integration errors and to prevent divergence near the axis where *r* → 0.

Once convergence is achieved, the total bending energies are evaluated by numerical integration of the energy density along the membrane contour. The results yield the total bending energy of the membrane as a function of the budding angle *α*, allowing for direct comparison between flat and vesicle geometries and identification of the most energetically favorable budding pathways. For the flat-membrane case, the same procedure is followed, except that the boundary conditions are modified from *r*(*s*_0_) = 0 and *ψ*(*s*_0_) = 0 to *ψ*^′^(*s*_0_) = 0 and *ψ*(*s*_0_) = *π*, consistent with our definition of *ψ*, in order to impose a flat membrane profile at the boundary. A detailed derivation for the flat-membrane case is provided in the Appendix. Here we focus on equilibrium membrane shapes and energetic trends; incorporating hydrodynamic dissipation and kinetic timescales would require a fully dynamical framework and is therefore left for future work.

### B. Transfection of cultured cells to produce viral particles

HEK-293T cells (ATTC) were grown in complete DMEM medium (Corning, Inc.) (1g/l glucose, 100 mg/L sodium pyruvate, and 10% certified fetal bovine serum (SigmaAldrich, Inc.)) at 37ºC and 5% CO2. Viral particles were produced by transfection using the plasmid pACNR-CHIKV described in Colunga-Saucedo *et al*.^38^. The transfections were performed in 6-well (Corning, Inc.) plates using Lipofectamine 2000 (Invitrogen, Inc.) according to the vendor’s protocol, with the modification that 1.8 *µ*g of plasmid was used to transfect 300,000 seeded cells. At 24 hours post-transfection (h.p.t.), the supernatant was removed and replaced with new complete DMEM, and at 48 h.p.t., the media was replaced with 2% glutaraldehyde in 0.1 M cacodylate buffer, pH 7.2, for 2 hours at room temperature, following a previously published protocol^39^. The resin discs were cut using a diamond blade on a LEICA EM UC7 ultramicrotome, placed on copper grids, and stained with 0.5% uranyl acetate and 0.5% lead citrate. The samples were visualized on a JEOL JEM-2100 transmission electron microscope at 200 kV with a 4k Gatan One View camera. The images were analyzed using ImageJ® (NIH)^40^ and Fiji (NIH) software^41^.

## III. RESULTS

To obtain the equilibrium membrane profiles at different stages of budding, characterized by varying values of the budding angle *α*, we numerically integrated the shape Eqs. (10)–(14). The resulting membrane profiles for budding through a vesicle and a flat membrane are shown in Figs. 3 and 4. Each contour describes the equilibrium configuration obtained by minimizing the total bending energy under the area constraint, with the cap area fraction *f* = 14%. Both the vesicle and flat geometries were constrained to have the same total area and cap size, ensuring that the geometry of the protein-covered cap is identical in both cases. This enables a direct comparison of their neck morphologies and associated energy profiles.

**FIG. 3.**
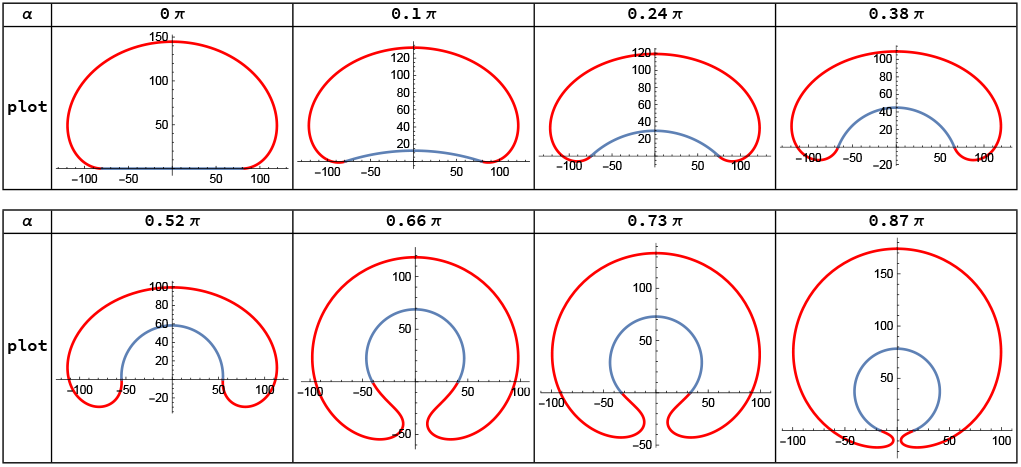
Membrane profiles at successive stages of budding from a vesicle-shaped membrane. Numerically computed profiles are shown for opening angles *α* = 0*π*, 0.1*π*, 0.24*π*, 0.38*π*, 0.52*π*, 0.66*π*, 0.73*π*, and 0.87*π* (from left to right, and top to bottom). The parameter *α* quantifies the extent of budding, with *α* = 0 representing a flat patch and *α* = *π* a fully closed vesicle. As *α* increases, the membrane curves inward, forming a neck that connects the protein-covered cap to the vesicle body. In this case, the preferred curvature is *R*_0_*/a* = 40 and the cap area fraction of the total area is *f* = 14%.

**FIG. 4.**
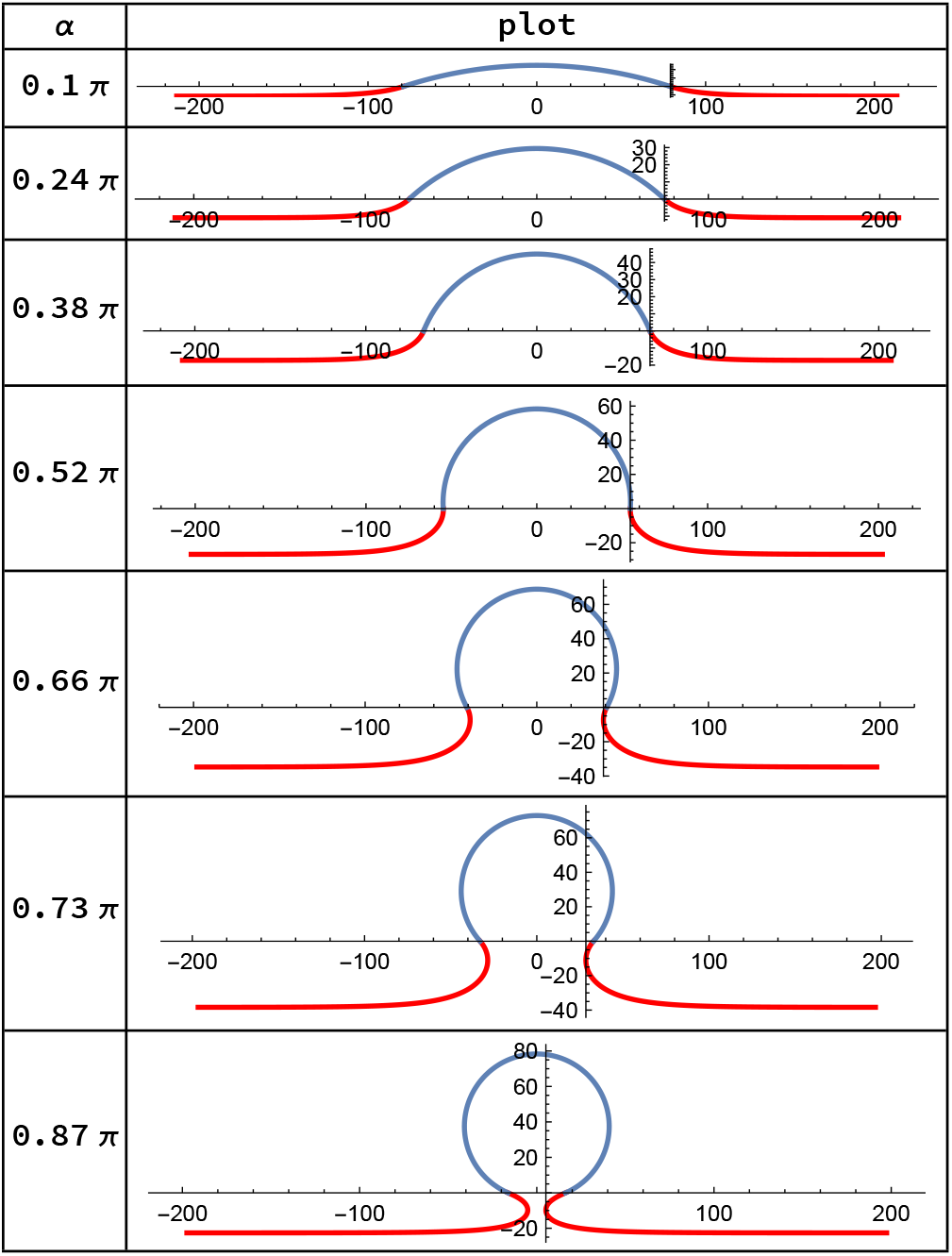
Membrane profiles at successive stages of budding from a flat membrane. Numerically computed membrane profiles are shown for the opening angles *α* = 0.1*π*, 0.24*π*, 0.38*π*, 0.52*π*, 0.66*π*, 0.73*π*, and 0.87*π* (from top to bottom). As in the vesicle case, increasing *α* leads to stronger membrane bending and the emergence of a narrow neck. Here, the preferred curvature is *R*_0_*/a* = 40, and the cap area fraction is *f* = 14%.

Fig. 3 displays the sequence of membrane profiles obtained for the vesicle geometry. The sequence illustrates the continuous morphological transition from a flat patch to an invaginated bud. As the budding angle *α* increases, the neck connecting the cap to the surrounding membrane gradually narrows, and the curvature near the junction becomes more pronounced. Near *α* ∼ *π*, the cap nearly encloses a complete sphere, corresponding to a fully formed bud ready for scission. Our modeling framework does not permit the scission step itself to occur.

Fig. 4 presents the corresponding results for the flat-membrane geometry. At the onset, the membrane remains entirely flat, representing an energetically favorable state. The emergence of a neck requires the cap region to locally deform the membrane and introduce curvature. The sequence illustrates the morphological evolution from a flat membrane serving as a reference for comparison with the vesicle geometry in Fig. 3.

Together, Figs. 3 and 4 show distinct budding pathways from flat and vesicular membranes under identical cap-to-membrane ratio and mechanical parameters. Both sequences highlight how the membrane deformation intensifies with increasing *α*, setting the stage for the subsequent energy analysis.

We note that in this paper we assume that the cap region has a preferred curvature, with *R*_0_*/a* = 40. Such curvature may originate from the intrinsic dihedral angles between protein subunits^42^, from RNA–protein interactions mediated by packaging signals^43^, from the radius of gyration of the genome considering RNA base-pairing and its stiffness^44^, or from curvature induced as a capsid buds through a membrane covered with spike proteins.

### A. Bending Energy Contributions from the Cap and Free Membrane

After determining the membrane profiles along the budding pathways for both vesicular and flat geometries, we next analyze the bending-energy contributions from the cap and the surrounding protein-free membrane region. For each geometry, the appropriate ground-state (reference) energy has been subtracted, and the analysis focuses on relative energy changes and energy gradients along the budding coordinate, which is related to the thermodynamic driving force for bud progression or stalling.

Because the cap area is constrained to be identical in both geometries and the spontaneous radius of curvature is fixed at *R*_0_, the bending energy of the cap is the same for the planar and vesicular cases and is given by Eq. 4, where *κ* is the bending rigidity, *R* the instantaneous radius of curvature of the cap, and *R*_0_ the radius of the fully closed spherical bud. The spontaneous curvature is set to 1*/R*_0_, representing the curvature induced by bound proteins or scaffolding complexes that promote budding.

As shown in Fig. 5(a), the cap energy varies identically with the budding angle *α* in both geometries; the curves for the flat membrane (green) and the vesicular membrane (red) completely overlap. All energies are expressed in units of the bending rigidity *κ*, measured in units of thermal energy *k*_B_*T* . As the value of *α* increases, the cap becomes more curved and the bending energy decreases monotonically. In the limit *α* → *π*, the shape approaches a spherical morphology of radius *R*_0_. Because scission is not included in the present model, however, the bud remains connected to the parent membrane through a narrow neck, which prevents the protein-covered region from attaining the exact spherical shape. As a result, the cap bending energy remains finite but becomes very small at large *α*. On the scale of Fig. 5(a), this residual energy is not resolved and therefore appears to approach zero.

**FIG. 5.**
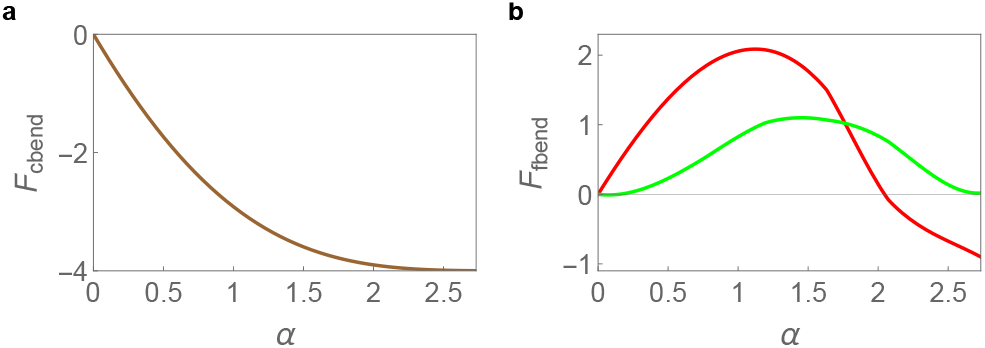
Cap and membrane bending energies *F*_cbend_(*α*) and *F*_fbend_(*α*) as a function of the polar angle *α*. Energies are shown relative to the corresponding ground-state reference for each geometry. (a) The bending energy decreases monotonically as the cap grows and the bud approaches closure. For a fixed cap area and spontaneous curvature, the bending energy is identical for the flat and vesicular geometries. The radius *R*_0_ is the spontaneous radius of the cap. (b) The membrane bending energy *F*_fbend_(*α*) is shown as a function of the opening angle *α* for the vesicle (red) and flat-membrane (green) geometries. In both cases, an energy barrier appears during the budding process. For the vesicle, the final energy lies below the initial energy, showing that bud completion is energetically favorable and curvature driven. In contrast, the flat-membrane geometry exhibits nearly identical initial and final energies, indicating a weak energetic driving toward closure and a higher likelihood of stalled budding.

The membrane energy is obtained from the solutions of Eqs. (10–14) for 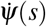, sin *ψ*(*s*), and *r*(*s*), which specify the local curvature and geometry of the membrane along the arc length *s*. By substituting these solutions into the bending energy expression (Eq. 5) and integrating from *s* = 0 to *s* = *s*_0_, we obtain the membrane energy for both the flat and vesicle cases.

As shown in Fig. 5(b), the membrane energy profiles exhibit energy barriers in both the flat and vesicle cases. As *α* increases, the vesicle budding case reaches its maximum energy barrier earlier (at a smaller opening angle) than the flat case, and the barrier height is also larger. This indicates that most of the membrane becomes flat again, except for a small and highly curved neck region, and the final energy remains slightly higher than the initial value. In contrast, for the vesicle case, the lower final energy indicates that bud completion is energetically favorable, providing a driving force for the process to proceed to full closure. This trend is also evident in the profile plots: when *α* is small, the membrane shape resembles an ellipse, whereas for large *α*, it becomes nearly spherical (see Fig. 3).

To obtain the total bending energy of the system, we combine the cap contribution with that of the surrounding free membrane as shown in Fig. 6(a). As the cap energy is identical for flat and vesicle membrane, any difference in energy arises solely from the deformation of the protein-free membrane region. For small *α*, the total bending energy reflects a competition between the decreasing cap bending energy and the increasing deformation energy of the surrounding membrane. As *α* increases, the total bending energy decreases monotonically. Again, all energies are evaluated relative to the corresponding ground-state reference configuration (planar membrane or spherical vesicle), and only energy differences along the budding pathway are physically meaningful. A physically relevant comparison is therefore obtained by examining the energy gradient, as shown in Fig. 6(b). Since the equilibrium state of the cap is that of a sphere, this eliminates the energy barrier in both cases, although the energy profiles differ as the budding process progresses.

**FIG. 6.**
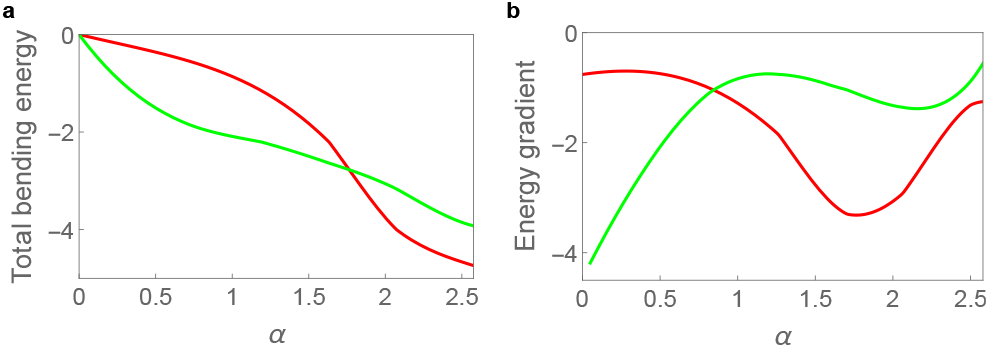
Total bending energy for membrane deformation in flat and vesicular geometries. (a) The plot shows the total bending energy as a function of the opening angle *α* for the vesicle (red) and flat-membrane (green) configurations. For small *α*, the total bending energy reflects the competing effects of decreasing cap bending energy and increasing membrane deformation energy. As *α* increases, the energy decreases monotonically in both cases. (b) The green and red curves show *dF/dα* versus *α* for the flat and vesicle geometries, respectively. The derivative *dF/dα* characterizes the energy gradient along the budding coordinate *α*.

Fig. 6(b) shows the derivative of the total bending energy with respect to the opening angle *α* for the vesicle (red) and flat-membrane (green) geometries. The derivative *dF/dα* represents the energy gradient along the budding coordinate *α*, which determines whether the system evolves toward or away from bud closure. Around *α* ≈ 1.4, the magnitude of this energy gradient is substantially larger for the vesicular geometry than for the flat membrane, indicating a stronger tendency toward bud completion for the vesicle case. This difference originates from the underlying membrane curvature, which alters the redistribution of bending stresses and facilitates neck formation. However, at early stages of budding, where *α* is small, the energy gradient is slightly larger for the flat membrane, suggesting that nucleation is more favorable on a planar surface before curvature effects become dominant.

The comparison between vesicle and flat case membrane energy profile highlights how pre-existing curvature in the vesicle geometry helps neck formation and completion of the budding process when comparing to the flat case.

### B. Line Tension

In our model, the cap is treated as a spherical segment. Because the cap area is fixed, its size and shape can be fully characterized by a single parameter, the opening angle *α*. Once *α* is specified, both the cap geometry and, consequently, the boundary circumference *L* are uniquely determined. Using the effective line tension parameter *λ* (defined in units of *κ/a*), the energetic cost associated with the boundary between the protein-covered cap and the surrounding protein-free membrane is incorporated as *E*_*line*_ = *λ L*. Since line tension acts only along this boundary, it does not influence the equilibrium membrane shape and is therefore excluded from the energy-minimization process. Its contribution is instead added to the total bending energy after the membrane shape is determined, allowing us to quantify its energetic effect.

The appropriate range of the line-tension coefficient *λ* depends on physical factors such as the density of protein aggregation and the specific viral system^4^. Comparison with SARS-CoV-2 experimental measurements suggests that effective line-tension values on the order of *λ* ≈ 0.005 (in units of *κ/a*) provide reasonable estimates for protein-associated viral membrane boundaries. However, *λ* may vary among different viruses depending on their lipid and protein composition^4,33,45–50^. Fig. 7(a) shows the influence of the line tension on the total bending energy of the system as a function of the opening angle *α*. The energy profiles for *λ* = 0, 0.005, and 1 are presented in Fig. 7(a). Solid, dashed, and dotted lines correspond to *λ* = 0, 0.005, and 1, respectively. As we do for all energies shown, for each case the reference energy has been subtracted so that the curves start from zero at *α* = 0. For the small value *λ* = 0.005, the curves nearly overlap with those for *λ* = 0, indicating that the contribution of line tension to the total energy is minimal. As *λ* increases to 1, the effect becomes more pronounced: the energy decreases relative to the *λ* = 0 case as *α* increases, reflecting the energetic contribution of line tension during budding.

**FIG. 7.**
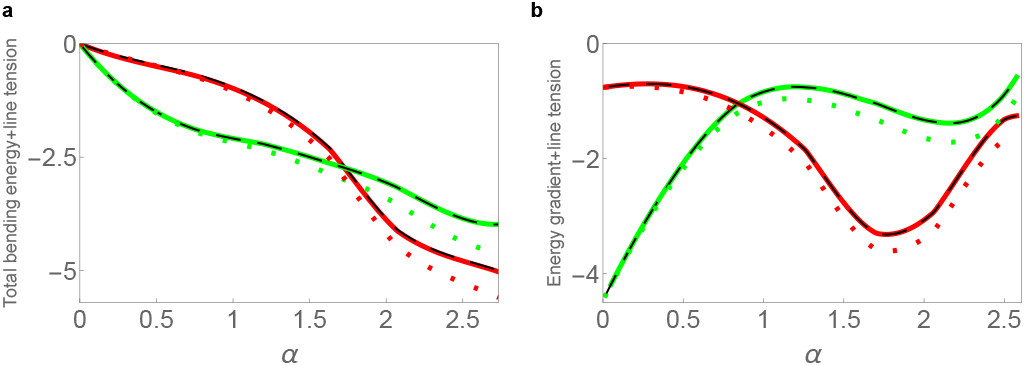
Total bending energy and energy gradient with line tension included. Total bending energy and its derivative with line tension included for different values of the line-tension parameter *λ* : *λ* = 0 (solid), *λ* = 0.005 (dashed, black), and *λ* = 1 (dotted), for both vesicular (red) and flat-membrane (green) geometries. (a) Because *λ* = 0.005 is small, the difference between *λ* = 0 and *λ* = 0.005 is not clearly visible. Increasing line tension (*λ* = 1) lowers the energy relative to *λ* = 0, with the difference increasing as *α* increases. (b) Energy gradient as a function of *α* for *λ* = 0, 0.005, and 1. The magnitude of the gradient increases with increasing *λ* .

The corresponding derivative of the total bending energy, including the line-tension contribution, is presented in Fig. 7(b). Because *λ* = 0.005 is small, the energy-gradient curves for *λ* = 0 and *λ* = 0.005 nearly overlap for each geometry. In contrast, for *λ* = 1 the overall trend remains similar to the *λ* = 0 case, but the magnitude of the energy gradient increases. This indicates that line tension strengthens the energetic driving toward bud closure by penalizing the presence of an open edge. Consequently, larger line tension reduces the stability of partially formed buds and favors complete vesicle formation.

### C. Experiments

In the previous section, we introduced a theoretical model describing the energetics of viral budding from both curved and flat membranes. To assess how well it captures the behavior of real viral particles, we performed experiments and generated TEM images of virus budding in infected cells. For this purpose, we used Chikungunya 181/25 virus (CHIKV), a prototypical alphavirus with a well-characterized assembly pathway. Alphavirus particles contain two concentric protein shells: an inner nucleocapsid (core), which packages the genomic RNA and forms an amorphous complex in the cytoplasm, and an outer shell composed of trimers of E1–E2 heterodimers embedded in the plasma membrane. Because the core binds to the cytoplasmic domains of these glycoproteins, assembly and budding proceed simultaneously at the plasma membrane^1^. In our model, we implicitly represent the core by prescribing a spherical cap. Since the resulting shapes can be uniformly scaled by the cap size, this allows for direct and quantitative comparison between our theoretical predictions and the experimental images.

Representative experimental images of CHIKV assembly and budding obtained by thin-section TEM are shown in Fig. 8. Briefly, CHIKV-infected cells were fixed, embedded in resin, and cut into ultrathin sections (approximately 50–80 nm). These sections were mounted on TEM grids, stained, and imaged using a transmission electron microscope (see Methods). This approach enables direct visualization of membrane-associated viral structures at high resolution.

**FIG. 8.**
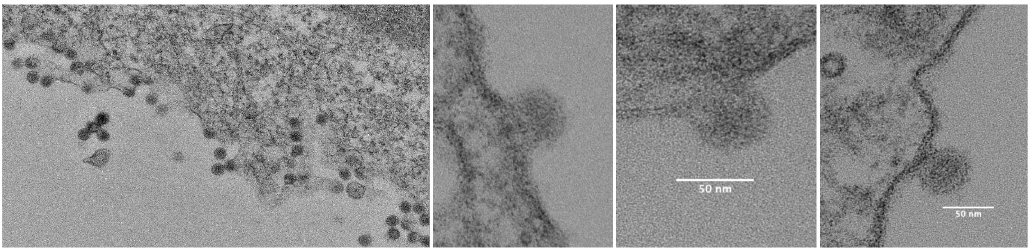
Thin-section TEM images of alphavirus budding from the plasma membrane of an infected cell. These images show distinct stages of the budding process and allow comparison with the theoretical shape contours derived from our model. Most budding caps appear as spherical segments, consistent with our geometrical assumption. The left panel shows multiple budding events, while the three panels on the right each display an individual budding profile.

From the resulting micrographs, we selected images that clearly capture full budding profiles of CHIKV particles at the host cell membrane. The left panel of Fig. 8 shows a low-magnification view containing virions in the extracellular space as well as multiple particles budding from the plasma membrane of a monolayer of cultured cells. The three images on the right, each with a scale bar of 50 nm, depict distinct stages of assembly and budding, along with the characteristic curvature of the plasma membrane.

## IV. DISCUSSION

Experimental images of HIV budding (Fig. 1) and alphaviruses (Fig. 8) show that budding rarely occurs on a perfectly flat plasma membrane. Owing to the intrinsic fluidity and deformability of lipid bilayers, local regions of the membrane often possess finite curvature even before budding begins. In many cases, budding is initiated in areas where the membrane is already slightly deformed, and neighboring budding events can further perturb the local geometry, effectively modifying the boundary conditions at the bud-membrane interface. These observations motivated us to investigate how boundary flexibility influences the energetics of budding.

In previous theoretical treatments, the far-field membrane was typically assumed to be flat 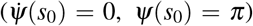. In the present study, we relaxed this condition by allowing 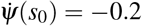 and *ψ*(*s*_0_) = 1.03*π*, representing a membrane that is not perfectly flat but exhibits a small finite slope at the outer boundary as shown in Fig. 9(a), consistent with experimentally observed local curvature. Our calculations show that when the boundary allows a finite 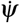 and a small deviation from *ψ*(*s*_0_) = *π*, the membrane develops a longer and smoother neck, and the total bending energy decreases more steadily as a function of *α* (Fig. 9(b)). In contrast, enforcing a strictly flat boundary produces a highly curved and short neck, leading to an energy profile with an extended plateau where the slope *dF/dα* is very small. This plateau corresponds to a regime where the energy gradient is small, leading to weak energetic driving toward closure and potentially hindering the completion of budding.

**FIG. 9.**
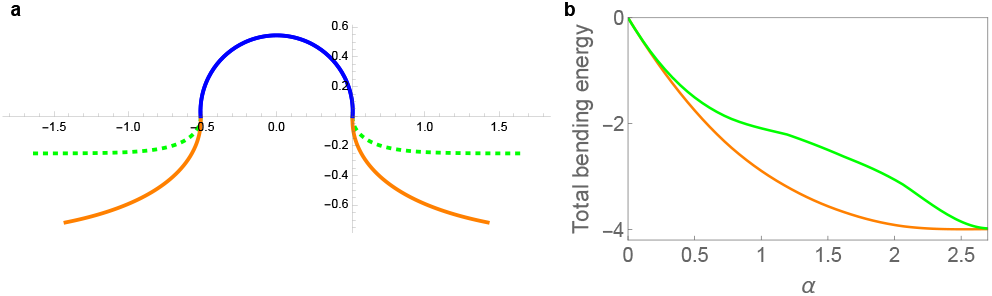
Boundary-condition effects on membrane shape and total bending energy. (a) For this configuration, the cap geometry is fixed by *α* = 0.52*π*, so all contours share the same spherical-cap shape. The dotted green curve corresponds to a membrane that remains flat far from the budding site 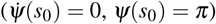. The solid orange curve shows the case where the far field is allowed a small slope and a slight deviation from flatness 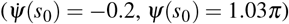. Allowing this deviation produces a longer, less curved neck and a larger budding height. Here, 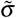 decreases from 26.5 (green) to 0.7 (orange). We emphasize that 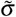 denotes a Lagrange multiplier enforcing global membrane area conservation, rather than a directly measurable physical membrane tension. The two contours share the same cap area fraction, *f* = 14%. (b) The orange curve shows the total bending energy as a function of *α* for the boundary condition 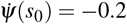, *ψ*(*s*_0_) = 1.03*π*. Compared with the energy profile for the strictly flat boundary (Fig. 6(a), green curve), the region between *α* = 1 and 2 no longer exhibits a plateau; instead, the energy decreases smoothly throughout the budding process.

In our model, the parameter 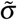 acts as a Lagrange multiplier that enforces the membrane area constraint. Although *σ* has the same dimensions as a physical surface tension, it represents a constraint-associated energy density and should not be compared directly with experimentally measured rupture thresholds. Consequently, 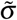 adjusts self-consistently during the budding process, effectively playing a role analogous to a chemical potential for membrane area. We find that the magnitude of 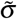 is substantially higher for the flat-membrane case than for the vesicular geometry. This increase arises from the geometric constraint imposed by the flat boundary, which resists curvature formation. Relaxing the boundary condition reduces the required constraint strength and lowers the associated bending-energy cost.

Taken together, these results suggest that local membrane curvature and boundary flexibility can facilitate bud completion. In a cellular context, nearby budding events or preexisting membrane curvature may therefore create a mechanically favorable environment for efficient viral budding.

### A. Quantitative Comparison of Theoretical Predictions with Experimental Observations of Alphavirus Budding

To demonstrate the predictive power of our theoretical framework, we quantitatively compared the predicted membrane shapes with thin-section TEM images of alphavirus budding. Experimental membrane contours were extracted using the Segment Anything Model (SAM)^51^, which is an AI-based segmentation framework that can extract objects from images with given prompts, followed by edge-detection analysis to reconstruct the bilayer geometry with sub-pixel precision. The resulting coordinates were directly compared with theoretical profiles obtained from numerical solutions of the shape equations.

Because the membrane in the experimental image is not perfectly flat, we compare the data with the theoretical shape obtained using the boundary conditions 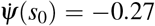 and *ψ*(*s*_0_) = *π*. Fig. 10 shows that the equilibrium shapes predicted by the model are consistent with experimentally observed budding morphologies. The TEM images correspond to near-circular cross sections of budding virions, which are expected when the sectioning plane passes close to the symmetry axis of an approximately axisymmetric bud. Overall, the theoretical contour captures the main geometric features of the experimental membrane profile. However, careful inspection reveals a small but systematic deviation on one side of the bud, where the experimental membrane contour exhibits slightly stronger curvature than predicted.

**FIG. 10.**
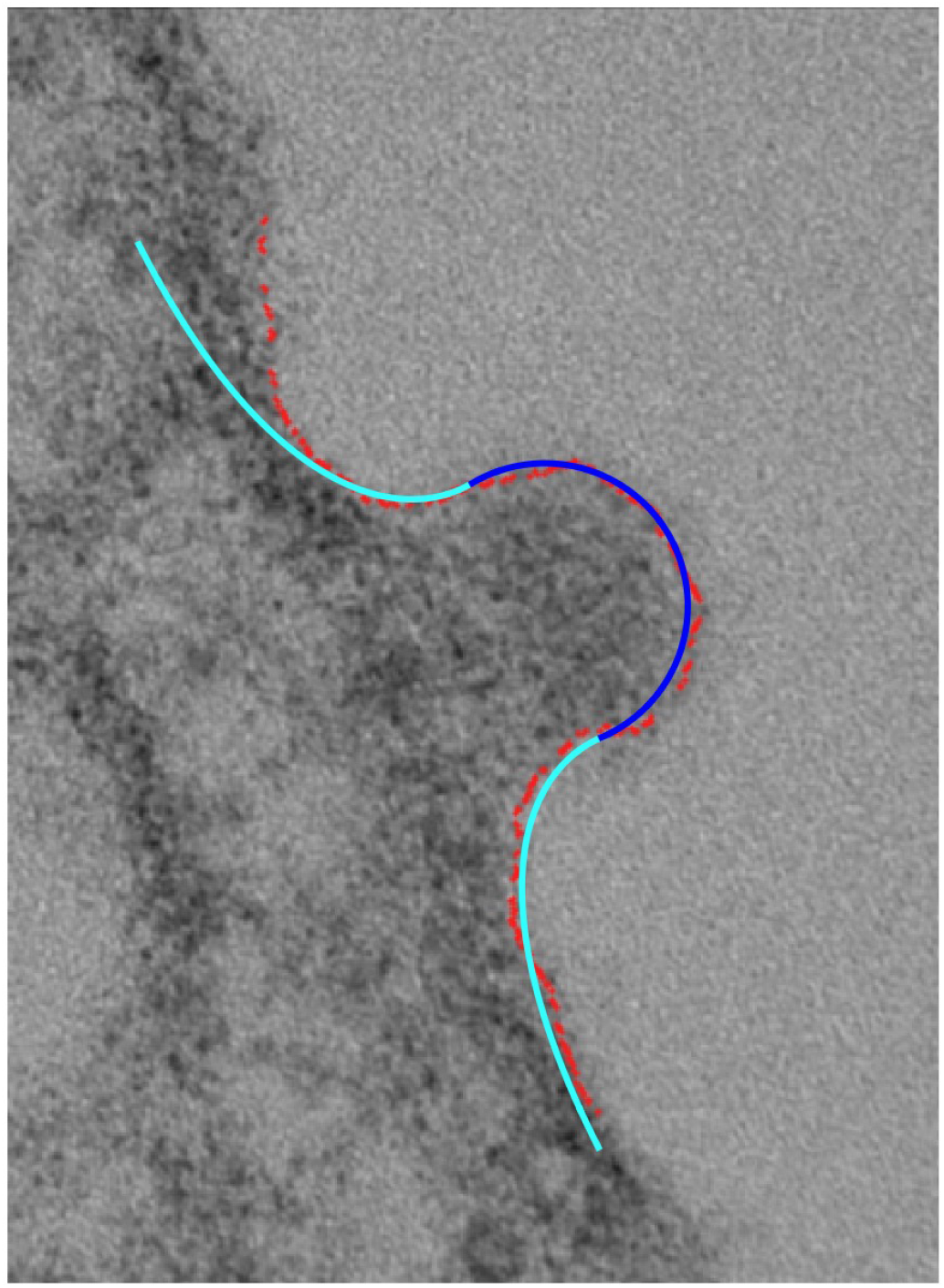
Thin-section TEM image of alphavirus budding and comparison with the theoretical profile under symmetric boundary conditions. The red dots show membrane contours extracted from the TEM image using the Segmentation Anything Model (SAM)^51^. The blue and cyan curves represent the protein-covered cap and the adjoining protein-free membrane predicted by the theoretical model, respectively. The boundary condition for membrane (cyan) is 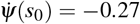 and *ψ*(*s*_0_) = *π*. The close agreement between the theoretical and experimental contours indicates that the model accurately reproduces most of the membrane profile, except for a small deviation on one side of the boundary observed during alphavirus budding.

Arguably, this discrepancy originates from asymmetry in the experimental membrane shape. In the TEM image, the left and right sides of the budding profile are not mirror-symmetric: one side of the membrane connects to the surrounding bilayer at a slightly steeper angle. Additional images of asymmetric budding are shown in Fig. 13 in the Supplementary Materials. Such asymmetry likely reflects local variations in lipid composition or interactions with adjacent budding sites, both of which can perturb the local curvature. To account for this feature, we modified the theoretical boundary condition to allow an asymmetric slope at the cap–membrane junction. Specifically, the values of 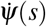 and *ψ*(*s*) were allowed to differ slightly on the two sides of the bud, producing a small angular offset between the left and right membrane branches. The asymmetric boundary condition is 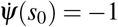 and *ψ*(*s*_0_) = 0.75*π*.

The resulting asymmetric solution, shown in Fig. 11, provides a significantly improved match to the experimental contour. The theoretical profile now reproduces not only the cap curvature and neck width but also the subtle difference in slope between the two sides of the membrane. This agreement confirms that the asymmetry observed in the TEM image is intrinsic and not due to noise or image-processing artifacts. Physically, the result highlights how even small differences in local boundary curvature can alter the overall membrane shape, producing the asymmetric budding profiles often seen in cryo-EM and TEM studies of alphavirus and other enveloped viruses.

**FIG. 11.**
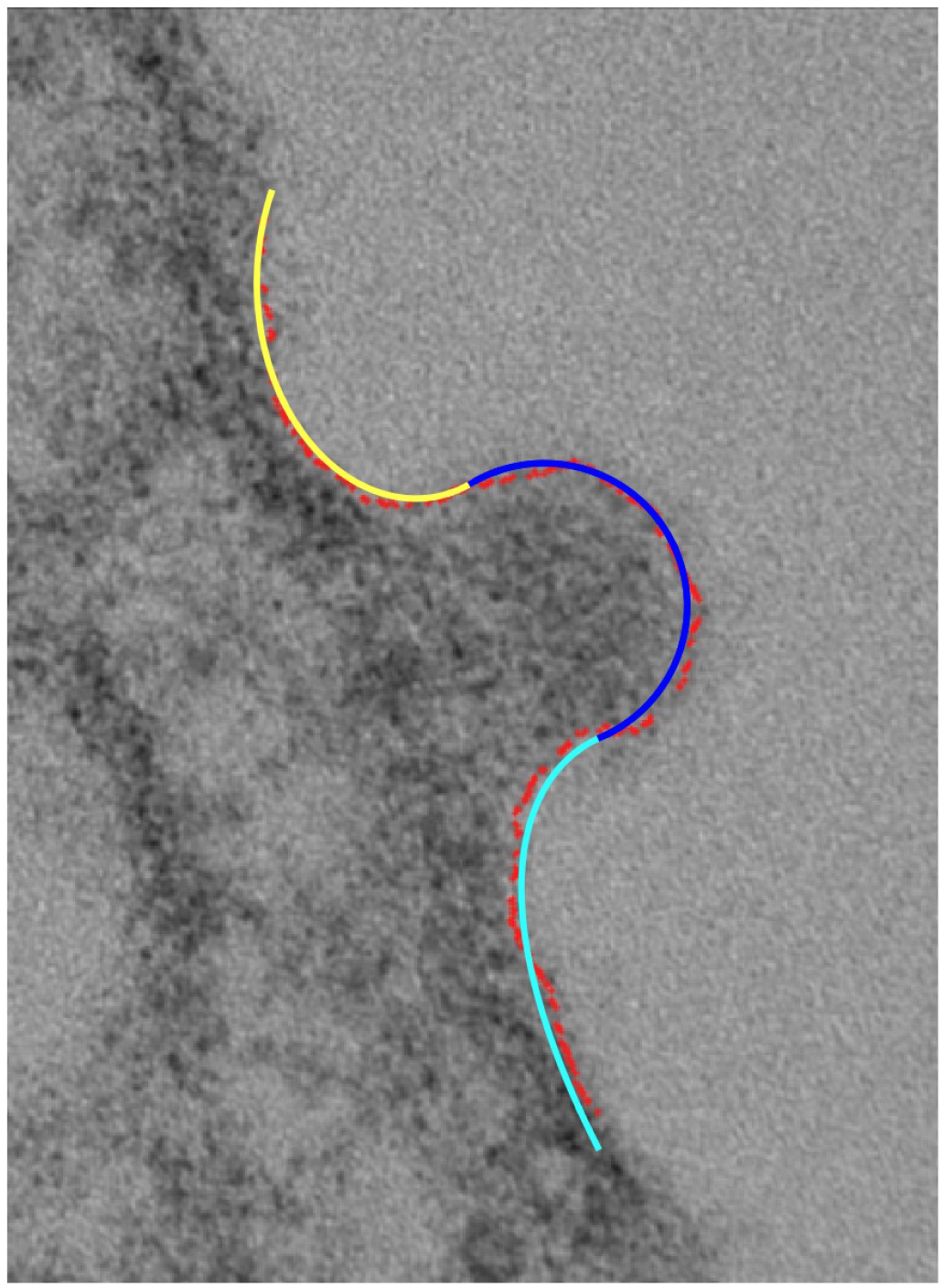
Thin-section TEM image of alphavirus budding and comparison with the theoretical membrane profile under asymmetric boundary conditions. The experimental membrane contour (red dots) was extracted from the TEM image using the Segmentation Anything Model (SAM). The blue and cyan curves represent the theoretical cap and adjoining protein-free membrane regions, respectively, while the yellow curve shows the theoretical profile obtained with a modified, asymmetric boundary condition. Introducing this asymmetric boundary condition significantly improves the fit to the experimental membrane shape, particularly on the left side of the bud, where local curvature differs from that of a symmetric configuration. The asymmetric boundary condition is 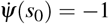 and *ψ*(*s*_0_) = 0.75*π*.

These findings emphasize that the perfectly flat and symmetric boundary conditions commonly assumed in theoretical treatments do not always reflect the true cellular environment. In cells, viral budding frequently occurs in regions where the membrane is already distorted by neighboring buds or scaffold proteins, naturally leading to asymmetric curvature at the boundaries. Our calculations show that relaxing both flatness and symmetry, allowing the far-field membrane to deviate slightly from a perfectly planar and mirror-symmetric configuration, markedly reduces the energetic cost associated with membrane deformation and smooths the energy profile as a function of *α*. This reduction in energetic cost lowers the energy barrier for bud completion, providing a physical explanation for the experimentally observed tendency of viruses to bud adjacent to one another.

These results connect directly to our comparison between vesicle and flat-membrane geometries. Asymmetric boundary conditions yield energy profiles that closely resemble those of the vesicle case, characterized by smoother deformation pathways and enhanced budding efficiency. Taken together, the theoretical and experimental analyses demonstrate that incorporating boundary asymmetry is essential for reproducing experimentally observed geometries and for understanding how local curvature, boundary flexibility, and asymmetry collectively promote cooperative viral budding.

The analysis presented above provides a general physical perspective for understanding membrane remodeling in viral systems, including cryo-electron tomography observations of SARS-CoV-2 virions that reveal the in situ architecture of viral envelopes and intermediate stages of membrane deformation during viral fusion^52^, a process that will be investigated in future work.

## V. CONCLUSION

Our theoretical and experimental analyses together reveal how membrane geometry and boundary conditions regulate the energetics and completion of viral budding. The energetic pathways for budding from a vesicle and from a flat membrane are fundamentally distinct. In vesicle-like geometries, such as the ERGIC where SARS-CoV-2 assembles, the underlying curvature initially raises the energy but produces a steeper energy gradient along the budding coordinate once budding begins, thereby favoring rapid and complete closure. In contrast, budding from a flat membrane begins in a low-energy state but encounters an extended plateau in the energy profile, corresponding to a weak energy gradient that can stall bud completion.

Relaxing the far-field boundary condition of the flat membrane, allowing small deviations from perfect flatness, makes the energy profile smoother and more similar to that of the vesicle case. Both the bending energy and the overall energetic cost of deformation decrease, explaining why viruses frequently bud adjacent to one another or within locally curved regions. Neighboring buds effectively soften the boundary, reducing the energy barrier for subsequent budding and promoting cooperative, curvature-assisted assembly.

Comparison between theory and thin-section TEM images of alphavirus budding demonstrates that the equilibrium shapes predicted by the model are consistent with experimentally observed membrane morphologies. The theoretical membrane contours capture key geometric features of the budding profiles, including overall curvature and neck morphology across different stages of budding. This level of agreement supports the physical framework of the model and illustrates how membrane geometry and boundary conditions shape the budding process.

Overall, these results establish a unified physical picture of viral budding in which local membrane curvature, boundary flexibility, and geometric coupling determine the energy gradients along the budding pathway that control initiation and completion. Curved or vesicle-like environments favor efficient closure, while boundary relaxation and cooperative budding on flat membranes provide alternative routes for overcoming energetic barriers.

## AUTHOR CONTRIBUTIONS

**Shuyang Zhang:** Data curation(equal), Investigation(equal), Formal analysis(equal), Methodology(equal), Visualization(equal), Writing – original draft(equal), Writing review and editing(equal). **Siyu Li:** Conceptualization(equal), Formal analysis(equal), Methodology(equal). **Miguel Angel Coronado-Ipiña:** Visualization(equal). **Mauricio Comas-Garcia:** Investigation(equal), Visualization(equal), Writing – review and editing(equal). **Ajay Gopinathan:** Conceptualization(equal), Investigation(equal), Formal analysis(equal), Methodology(equal), Writing – review and editing(equal). **Paul van der Schoot:** Conceptualization(equal), Investigation(equal), Formal analysis(equal), Methodology(equal), Writing – review and editing(equal). **Roya Zandi:** Conceptualization(equal), Data curation(equal), Investigation(equal), Formal analysis(equal), Funding acquisition(equal), Methodology(equal), Supervision(equal), Visualization(equal), Writing – original draft(equal), Writing review and editing(equal).

## CONFLICTS OF INTEREST

The authors have no conflicts to disclose.

## DATA AVAILABILITY

All theoretical results and numerical values supporting this study are contained within the paper. Experimental images used for comparison are included within the article and originate from collaborating laboratories or previously published sources as cited. No additional datasets were generated.

## ACKNOWLEDGEMENTS

This work was supported by University of California Office of the President UC Multicampus Research Programs and Initiatives, grant M21PR3267 (A.G.,R.Z.); the National Science Foundation under Grant No. NSF DMR-2131963 and Grant No. MCB/PHY-2413062 (RZ); the NSF-CREST: Center for Cellular and Biomolecular Machines at UC Merced (NSF-HRD-2112675 to AG) and the NSF Center for Engineering Mechanobiology grant (NSF-CMMI-1548571 subaward to A.G.); M.A.C.I. (CVU: 1232434) acknowledges the support of SECIHTI through the Ph.D. fellowship 4044561 M.C-G. thanks SECIHTI for the grant CBF2023-2024-1125 and COPOCYT for the grant 2024-03-M07 and the CIC-SaB/UASLP for Intramural Funding.

## Appendix A Derivation for the flat case

For the flat case, we also use *ψ*(*s*) to represent the shape and connect it with *r*(*s*) and *h*(*s*) as

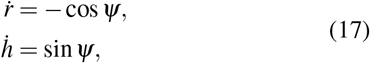

where the dot denotes differentiation with respect to the arc length *s*.

**FIG. 12.**
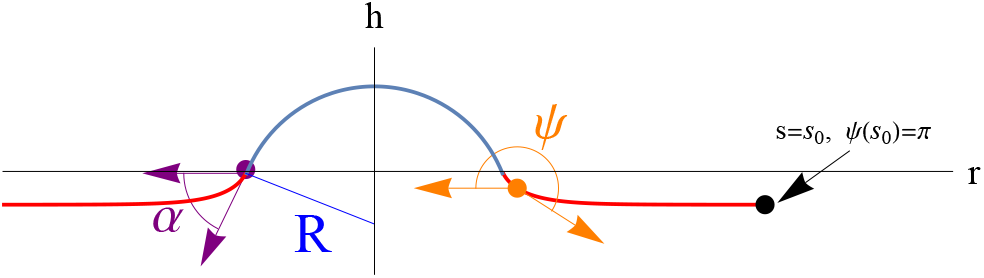
Schematic illustration of the budding geometry for the flat case.

The corresponding energy expressions are

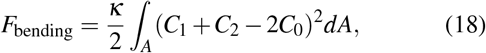

and

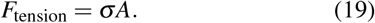

The membrane bending energy is described by the Helfrich functional, with a surface “tension” term enforcing the constraint of fixed total area. For a given *α*, the cap geometry is prescribed, and the associated line-tension contribution does not enter the shape determination and is therefore omitted here.

The total bending energy consists of contributions from the protein-coated cap and the surrounding bare membrane. The cap is modeled as a rigid spherical segment with analytically evaluated bending energy, while the elastic contribution of the protein-free membrane is obtained numerically from the membrane shape equations.

The cap bending energy is

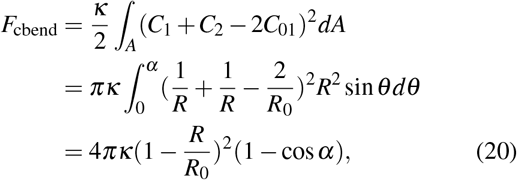

Here, 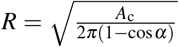 and is the radius of curvature of the cap and *A*_*c*_ *= f A* represents the area of the cap, where *f* is the fraction of the total area *A*. Here *θ* denotes the polar angle in spherical coordinates. The principal curvatures in cap area are *C*_1_ = *C*_2_ = 1*/R*. The spontaneous curvature of the cap is given by *C*_01_ = 1*/R*_0_ and 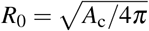.

The membrane bending energy *F*_fbend_ is

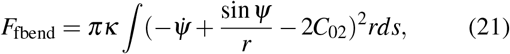

and the spontaneous curvature for membrane region is *C*_02_ = 0. Using the rotational symmetric arc-length parametrization, the two principal curvatures are given by (sin *ψ*)*/r* and 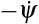, where, as before, the dot denotes differentiation with respect to the arc length *s*.

Followed by the energy functional 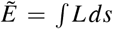, the Lagrangian *L* given by

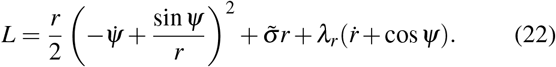

*λ*_*r*_(*s*) is a Lagrange multiplier and 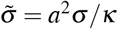 is also used in this derivation.

Since the Lagrangian does not depend explicitly on the arc length parameter *s*, the corresponding Hamiltonian is conserved. We therefore adopt a Hamiltonian formulation, which also allows the governing equations to be written as a set of first-order differential equations suitable for numerical integration.

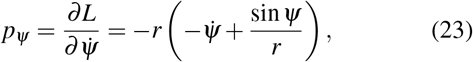

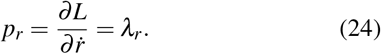

The Hamiltonian is

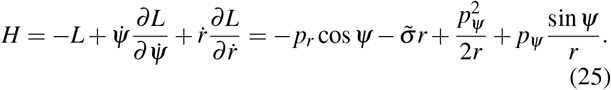

Introducing the conjugate momenta *p*_*ψ*_ and *p*_*r*_, the Legendre transform yields the Hamiltonian *H*, from which the equilibrium membrane shape is obtained through the associated Hamilton equations. This results in the first-order differential equations for the variables (*r, h, ψ, p*_*ψ*_, *p*_*r*_), which are solved numerically.

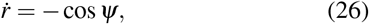

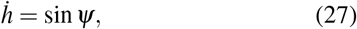

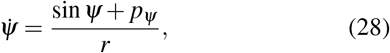

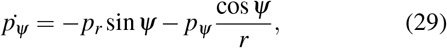

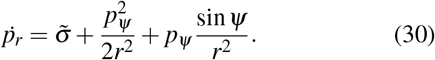

The equations are supplemented by boundary conditions. At the interface between the protein-covered cap and the free membrane (*s* = 0), continuity requires

- *r*(0) = *R* sin *α*;
- *h*(0) = 0;
- *ψ*(0) = *π* + *α*.

The initial slope 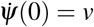 is treated as a shooting parameter. The corresponding initial values of the conjugate momenta *p*_*ψ*_ (0) and *p*_*r*_(0) are obtained from the Hamiltonian relations and boundary conditions.

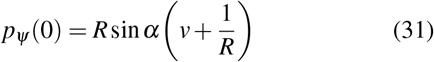

and

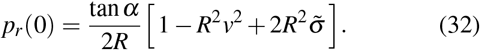

To ensure the flat membrane at *s* = *s*_0_, the following boundary conditions are imposed:

- 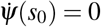, requiring the membrane curvature to vanish at the boundary, which ensures a smooth, flat continuation of the membrane contour;
- *ψ*(*s*_0_) = *π*, prescribing a horizontal membrane orientation at the boundary.

The resulting system of first-order differential equations is solved numerically using a fourth-order Runge–Kutta method. The initial slope 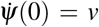 is tuned iteratively so that all boundary conditions are satisfied. Integration is performed from the cap–membrane junction(*s* = 0) toward the membrane edge(*s* = *s*_0_), and convergence is achieved when *ψ*(*s*_0_) = *π* and 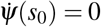 are met within a tolerance of 0.005.

## Appendix B Experimental Images

As shown in Fig. 13(a)-(c), numerous characteristic budding events are observed in these thin-section TEM images. Most budding events occur in proximity to neighboring structures or on membrane surfaces that are not perfectly flat, indicating that the budding process is often asymmetric.

**FIG. 13.**
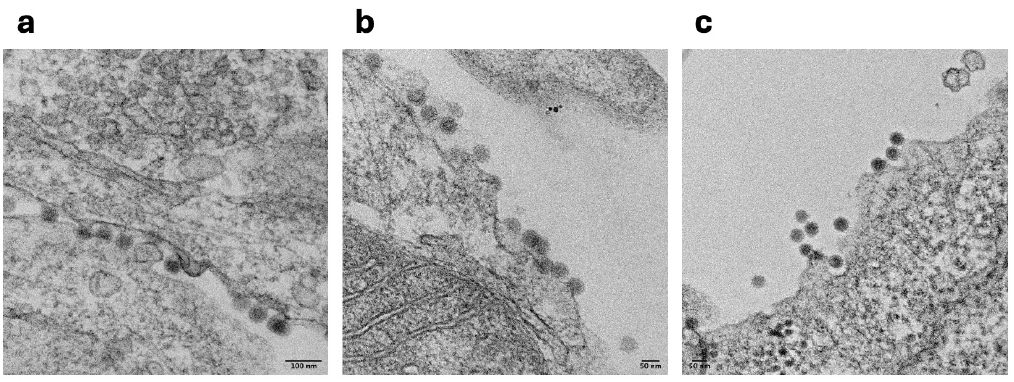
Experimental Images of asymmetric Chikungunya virus budding.

## Notes

### Competing Interest Statement

The authors have declared no competing interest.

